# An autonomous tail-beating cycle period expressed by a region-specific swimming pattern generator in the *Ciona* larva

**DOI:** 10.1101/2021.04.12.439438

**Authors:** Takashi Hara, Shuya Hasegawa, Yasushi Iwatani, Atsuo S. Nishino

## Abstract

Swimming locomotion in aquatic vertebrates, such as fish and tadpoles, is expressed through orchestrated operations of central pattern generators. These parallel neuronal circuits are ubiquitously distributed and mutually coupled along the spinal cord to express undulation patterns accommodated to efferent and afferent inputs. While such sets of schemes have been shown in vertebrates, the evolutionary origin of those mechanisms along the chordate phylogeny remains unclear. Ascidians, representing a sister group of vertebrates, give rise to tadpole larvae that freely swim in seawater. In this study, we tried to locate the swimming pattern generator in larvae of the ascidian *Ciona* by examining locomotor ability of segmented body fragments. Our experiments demonstrated necessary and sufficient pattern generator activity in a short region (∼10% of the body length as the longest estimation) including the trunk-tail junction but excluding most of the trunk and tail with major sensory apparatuses therein. Moreover, we found that these “mid-piece” body fragments express periodic tail beating bursts with ∼20-s intervals without any exogenous stimuli. Comparisons among temporal patterns of tail beating bursts expressed by the mid-piece fragments and by whole larvae placed under different sensory conditions suggested that the presence of parts other than the critical mid-piece had effects to shorten swimming burst intervals, especially in the dark, and also to expand the variance in burst durations. We propose that *Ciona* larvae perform swimming as modified representations of autonomous and periodic pattern generator drives, which operate locally in the region of the trunk-tail junction.

**Summary statement:** Mid-piece fragments of tadpole larvae of the ascidian *Ciona*, lacking most of the anterior trunk and posterior tail, autonomously and periodically express tail beating bursts.

## INTRODUCTION

Ascidians in general give rise to simplified tadpole-shaped larvae, which represent a miniature form of chordate bodyplans (e.g., Satoh, 2003; Meinertzhagen et al., 2004). Ascidian urochordates constitute a group in the phylum Chordata [or the superphylum Chordata (Satoh et al., 2014)] and are included in the sister clade of vertebrates. While adult ascidians are sessile and benthic, their tadpole larvae exhibit sophisticated swimming performance (Crisp and Ghobashy, 1971; Svane and Young, 1989; McHenry and Patek, 2004; Nishino et al., 2011; Salas et al., 2018). They freely swim in seawater as “micromachines” to disperse their packages of genetic contents and to find a suitable place for settling and growing until they become adults (Hickman, 1999). The simplified architecture of larvae, composed only of thousands of cells, shares basic bodyplans with those of vertebrates, including the axial notochord, dorsal neural tube, and bilateral muscle bands in the tail (Kowalevsky, 1866; Katz, 1983). In the larva of *Ciona*, a representative model ascidian species, 18 electrically-coupled muscle cells are present on each of the left and right sides in the tail, the three-row arrangement (i.e. dorsal, medial, and ventral rows) of which is bilaterally mirror-imaged and invariant among individuals (Bone, 1992; Passamanech et al., 2007; Nishino et al., 2011; Razy-Krajka and Stolfi, 2019). In the larval central nervous system (CNS), ∼180 neurons have been identified, and the numbers, positions, and connections of their subtypes are also considered to be mostly invariable among individuals (Ryan et al., 2016; Hudson, 2016; Nishino et al., 2018). According to connectomic analysis by Ryan et al. (2016), 143 neurons are included in an anterior ganglion in the trunk, called the brain vesicle (BV), which contains a gravity sensor (called the otolith), a photoreceptor (ocellus), and their associating pigment cells. Twenty-five neurons are found in another ganglion, called the motor ganglion (MG), that is positioned close to the trunk-tail junction, whereas nine or more neurons are distributed along the caudal nerve cord (CNC) that runs in the tail (Ryan et al., 2016, 2017). Among these, five pairs of neurons in the MG that have synaptic contact with muscle cells have been identified as motor neurons (Stolfi and Levine, 2011; Nishino et al., 2011; Ryan et al., 2016, 2017; Nishino, 2018). These motor neurons extend posteriorly their axon along the CNC and innervate the dorsal muscle cells, except one pair of them innervating the anterior-most medial muscle cell as well (Nishino et al., 2011; Ryan et al., 2016, 2017; Nishino, 2018). Also, two pairs of ascending contralateral inhibitory neurons (ACINs) located in the anterior part of the CNC have been proposed to function for reciprocal tail beating (Horie et al., 2010; Nishino et al., 2010; Nishitsuji et al., 2012), although it has remained to be proved whether they really compose a necessary and sufficient element for swimming pattern generation.

Regulatory mechanisms for locomotion, such as swimming, walking, or flying, have been rigorously studied in vertebrate model systems. Those studies have illustrated that systemic components to form locomotion patterns by orchestrating musculatures are sufficiently equipped in their spinal cord and that qualitative and quantitative controls for driving the system (including gait choice) are dependent on efferent inputs from superior centers in the brain and afferent inputs from peripheral sensory systems (Hennemann, 1957; Grillner, 1975; Shik and Orlovsky, 1976; Fetcho and McLean, 2010; Grillner and El Manira, 2020). The neuronal systems in the spinal cord can work to express locomotion even without any afferent inputs from the periphery and thus are regarded to fulfill the definition of central pattern generator (CPG) (Bässler, 1986). Assuming apomorphic acquisitions of paired fins, legs, or wings in vertebrates, the ancestral means for locomotion in vertebrates would be axial undulation of the body, as exhibited by basal vertebrates, such as hagfish and lamprey, and non-vertebrate chordates, such as amphioxus and ascidian larvae (Young, 1950; Grillner, 1975). Operational mechanisms of the CPG for swimming have been extensively analyzed in spinal cords of lamprey, teleosts, and amphibian larvae (for reviews, Grillner et al., 1991; Roberts, 2000; Fetcho and McLean, 2010). It has been revealed so far that the undulatory swimming CPG is composed of neural circuits that are essentially composed of glutamatergic excitatory interneurons to positively regulate ipsilateral motor neurons and glycinergic inhibitory interneurons to negatively regulate contralateral inter- and motor neurons (e.g., Buchanan and Grillner, 1987; Grillner, 2003). It is also known that the basic circuits for the CPG are not localized but are ubiquitously distributed and mutually coupled in the spinal cord (e.g., Cangiano and Grillner, 2003; Wiggin et al., 2012). On the other hand, these CPG circuits do not operate in autonomous manners but are driven in response to efferent inputs from superior brain centers (e.g., reticular formation and midbrain locomotor region) or afferent inputs from cutaneous sensory system or by application of excitatory amino acids and their analogs, such as D- or L-glutamic acid, D-aspartate, L-cysteate, L-DOPA, and N-methyl-D-aspartate (Poon, 1980; Grillner and Wallén, 1984; Dale and Roberts, 1984; Grillner et al., 1991; Roberts et al., 2012). It is also intriguing that “fictive” swimming bursts emerge periodically and are separated by intervals, even when the spinal cords of lamprey or teleost fish are tonically stimulated by chemical activators (Cangiano and Grillner, 2003; Wiggin et al., 2012).

Considering these findings, we here tried to prove whether a swimming pattern generator (PG) of *Ciona* larva, if it exists, is ubiquitously distributed along the tail or localized to a certain region by examining locomotor ability in segmented parts of the larval body, approximately 1 mm long as a whole. Our results demonstrated a necessary and sufficient role of the MG and ACINs, supporting a localized swimming PG in *Ciona* larvae. We also found that “mid-piece” body fragments that lack body regions anterior to the MG and posterior half or more of the tail “swim” in an autonomous manner, i.e., without any exogeneous stimulators, highly periodically with ∼20-s intervals between tail beating bursts. We also examined temporal patterns of swimming behaviors of whole larvae with their displacement limited using methylcellulose seawater. Based on the results, we concluded that swimming patterns of the *Ciona* larva are expressed as modified representations by afferent and efferent regulatory inputs into the autonomously and periodically-driven PG that works in the region of the trunk-tail junction.

## MATERIALS AND METHODS

### Animals

Mature adults of *Ciona intestinalis* (Linnaeus, 1767) (called type A), which Brunetti et al. (2015) recently defined as *C. robusta* Hoshino and Tokioka, 1967, were obtained from populations reared in the Misaki Marine Biological Station, The University of Tokyo (Miura, Japan) or Maizuru Fisheries Research Station, Kyoto University (Maizuru, Japan) via National BioResource Project (NBRP), AMED, Japan. They were kept in 5 L laboratory tanks containing an artificial seawater (ASW; Marine Art BR, Osaka-Yakken, Japan) or a mixture of ASW with natural seawater. Seawater in the tanks was gently stirred using a paddle connected to a synchronous motor (15 rpm, Nidec Servo, Japan). The tanks were placed in an air-conditioned room (18-19℃) and kept under constant light to prevent uncontrolled releasing of gametes. Ascidians are hermaphroditic, and mature eggs and sperm obtained from different adults were cross-fertilized as described previously (Jokura et al., 2020). The fertilized eggs were reared in a cool incubator set at 18℃. Tadpole-shaped larvae hatched out at approximately 17-18 hours post-fertilization (hpf).

### Preparation of anterior and posterior fragments of *Ciona* larvae

Hatched larvae with a normal looking well-elongated tail (≤ 24 hpf) were selected and placed in ASW in a petri dish coated with silicone rubber (SYLGARD 184, Dow Corning). If required, a larva to be cut was pictured in advance with a digital camera (DiFi2, Nikon) or a video camera (HDR-CX420, Sony) mounted on a stereomicroscope (SMZ745T, Nikon) or with a digital high-speed camera (VCC-1000, Digimo, Japan) mounted on another stereomicroscope (MZ FLIII, Leica). When the larva ceased swimming, the larval body was cut into anterior and posterior fragments using vannas micro scissors (Type 501790, World Precision Instruments) at an arbitrary position. After the bisection, the anterior and posterior fragments were pictured again using the same photosystem, and the lengths of the total body, anterior fragment and posterior fragment were measured on a PC later to determine the position of the cut site using Bohboh image analyzer (ver. 3, Bohboh Software, Japan) or Image J (ver. 1.52a or 1.53c). Cut sites are represented herein as the relative location (%) on a scale set along the longitudinal axis of the body, in which the position of the trunk-tail junction was defined as 0% and that of the tail tip (excluding the larval tunic) was defined as 100%. Namely, when cut sites were in the trunk, they were represented by negative values (see Results). The values for cut sites were calculated by dividing the length of the remaining tail (when the cut site was in the tail) or of the remaining trunk (when the cut site was in the trunk) by the tail length before amputation. In the course of data analyses, we noticed that the sum of lengths of the anterior and posterior fragments after the amputation became significantly shorter than the sum of lengths of the trunk and the tail before cutting [94.9 ± 3.1%, mean ± standard deviation (s.d.), n = 125; samples represented in Fig. 1]. It is notable, therefore, that our estimation of cut-sites herein includes a tendency to show relatively lower values by about 5% (e.g., ∼95% for 100% or ∼9.5% for 10%). Considering other possible deformations of the fragments and/or variations of internal structures among the specimens (e.g., antero-posterior locations of particular neuron types), both of which were difficult to be evaluated, we simply show in the present report the values for the cut sites derived from the relative lengths of the remaining tail or trunk in the fragments.

**Fig. 1.**
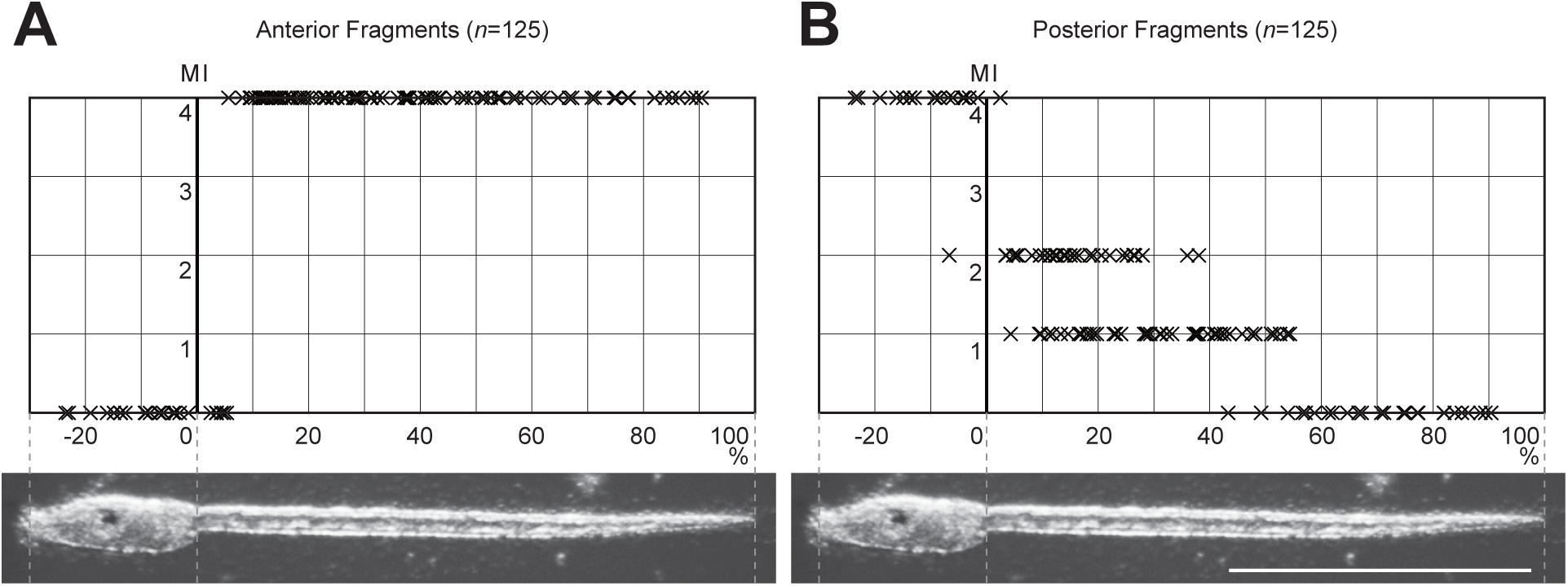
Locomotion ability of anterior and posterior fragments of segmented *Ciona* larvae. (A,B) Evaluated ability of movements of anterior (A) and posterior (B) fragments. The vertical axis indicates values in Motion Index (MI, see Table 1). Cut sites (marked by ×) are indicated along the horizontal axis, which is defined to be 0% at the trunk-tail junction and 100% at the tail tip. Pictures of a *Ciona* larva are shown at the bottom for reference. *n* denotes the number of fragments examined. Scale bar = 500 μm.

**Table 1.**
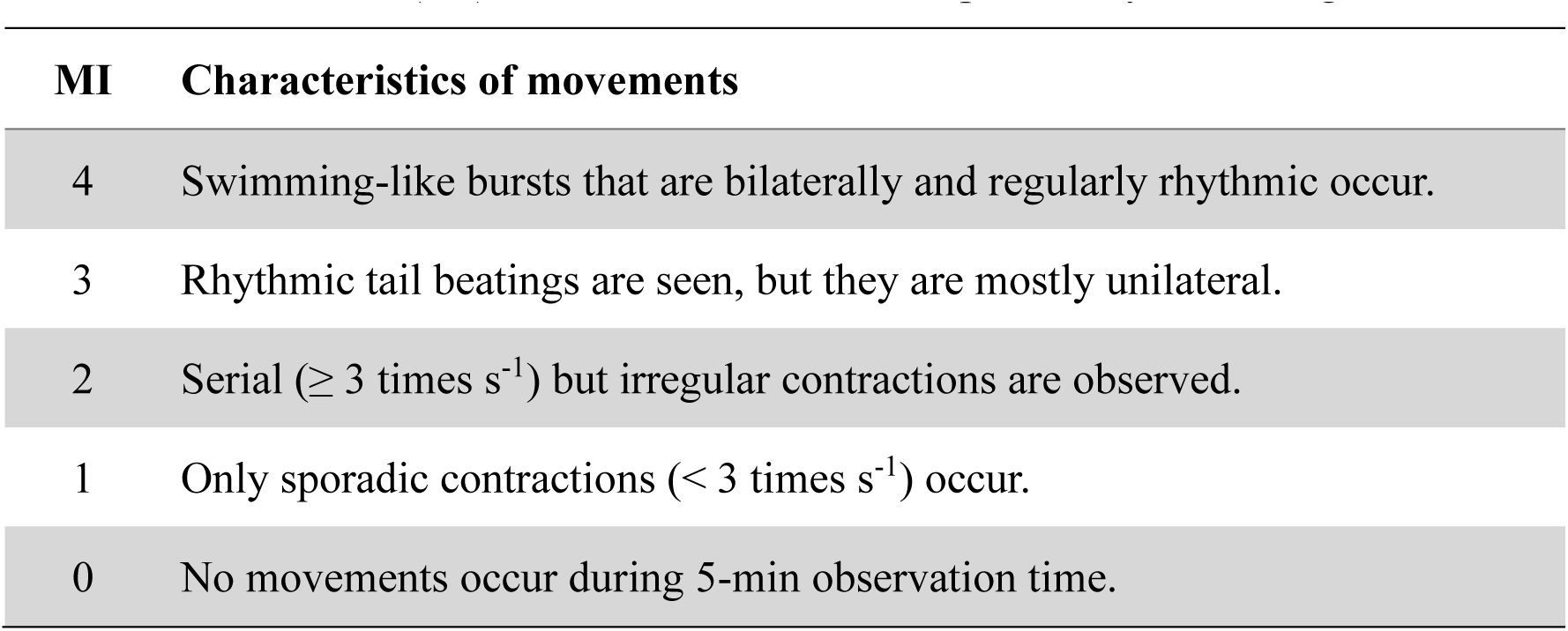
Motion index (MI) to evaluate movements expressed by larval fragments.

Both the anterior and posterior fragments were placed in another petri dish covered with silicone rubber containing ASW or ASW with 100 μM L-glutamic acid (L-Glu; Fujifilm Wako, Japan). After acclimation to the place and the solution for at least 20 min, movement patterns expressed by the anterior and posterior fragments were captured by a video camera or the high-speed camera (the same as above) mounted on the stereomicroscope, and the movement patterns were evaluated by direct observation or observation by the movies. We categorized the movement patterns into five levels, 0 to 4, and herein call them “motion index (MI)” values (Table 1). When tail beatings in a swimming burst were bilaterally and regularly rhythmic (namely “well patterned”), we evaluated the ability of the fragment as MI = 4. When tail beatings were rhythmic but unilateral, the ability was categorized as MI = 3. If burst-like serial contractions (≥ 3 times s^-1^) were observed, but their timings were irregular, motion ability of the fragment was defined as MI = 2. When only sporadic contractions (< 3 times s-1) occurred, the fragment was categorized as MI = 1. When we did not see any movements for 5 min, we defined MI of the fragment as 0 (Table 1). Evaluated MI values of the anterior and posterior fragments were plotted against the cut sites, respectively.

### Preparation of “mid-piece” fragments

“Mid-piece” fragments were defined as the body part from which the anterior trunk including the BV and the posterior half (or more) of the tail were both excluded but which retained MI = 4. To prepare such fragments, the posterior half of the tail of hatched larvae (≤ 24 hpf) was first cut off and then the anterior portion of the trunk was amputated so as to exclude the BV using the same tools as those described above. The location of the BV was identified by the pigmentations in the otolith and the ocellus (Katz, 1983), and we intended to sufficiently exclude the posterior BV region, which is important for controlling motor behaviors (Kourakis et al., 2019). After we confirmed that their MI was 4, we placed the mid-piece fragments in ASW or ASW containing 100 μM L-Glu or 1 mM L-Glu. Movements of the mid-piece fragments were recorded for 20 min by the video camera mounted on the stereomicroscope. By observing the recorded movies, the number of events was counted and the cumulative duration of tail beating bursts was measured.

“Mid-piece” fragments were prepared from hatched larvae at 19-20 hpf or 22-24 hpf by the procedure described above and their movement patterns were analyzed in detail. Movements expressed by mid-piece fragments were recorded as described above for 20 min, and were analyzed later on a PC. Movie files recorded by a video camera (HDR- CX420, Sony or GZ-F270-W, JVC) were exported as sequential images at 30 frames s^-1^ using a locally developed application running on a Windows PC. The images were imported into Image J, and changes in 8-bit grayscale brightness (0-255) at a “point of interest (POI)” were visualized. When the larval tail beats on an image sequence, the shade of the tail traverses the POI as many times as the tail beats and thus the brightness at the POI changes according to the movement of the tail. We also used the high-speed video camera (VCC-1000, Digimo, Japan) to calculate tail beating frequency at a higher resolution of time (250 frames s^-1^). For evaluation of durations and intervals of tail beating bursts, sporadic contractions, including flick-like movements, were excluded from our consideration, and swimming-like bursts of regular bilateral tail beatings (well-patterned movements) were taken into account.

### Analyses of durations and intervals of tail beating bursts by whole larvae

We intended to examine how periodic the movements expressed by unfragmented whole larvae are. We utilized methylcellulose (MC) to increase the viscosity of the medium (ASW) for swimming whole larvae, considering that higher viscosity would limit the amount of displacement by swimming, and that it would restrict otherwise variable sensory inputs (accidental collision with the dish bottom/wall or water surface as well as changes in the body direction to illuminated light, which are associated with free swimming within a dish). A few hatched larvae (19-20 hpf) were transferred into ASW containing 0.5% or 1% MC and then the medium was mixed well. After allowing time for acclimation to the viscous media for 10 min, we recorded movements expressed by the larvae for 15-20 min using the stereomicroscope and video camera as described above. Again, “swimming” bursts with regular bilateral tail beatings (well-patterned movements) were taken into account for evaluating durations and intervals of bursts, and sporadic contractions of the tail, including flick-like movements, were ignored.

In order to estimate the artificial effect of increased viscosity of the water medium on expression of movements, we compared tail-beating frequencies in swimming bursts of whole larvae in ASW containing 0% (namely, ASW itself), 0.5%, 1%, or 2% MC. For calculation of tail-beating frequencies, we placed several larvae (∼24 hpf) in ASW or ASW containing 0.5%, 1%, or 2% MC and then mixed the solution well. Swimming-like bursts of tail beatings were recorded using the high-speed camera as described above (VCC-H1000, Digimo, Japan) mounted on a stereomicroscope (MZIII, Leica) at 250 frame s^-1^. Since we found that the first and the last beating cycles in a series of tail beats (a burst) occur often irregularly, we calculated the frequency of tail beatings from durations of five tail beating cycles excluding the first and the last ones. The light condition was kept constant among different conditions of MC. All the data obtained above were analyzed using Microsoft Excel and R (ver. 3.5.0). The movie files were prepared using ImageJ (ver. 1.53c), Video Editor on a Windows-10 PC, and iMovie on an iPad Pro.

## RESULTS

### Experimental identification of the localized swimming PG

Neural circuits for the swimming CPG are ubiquitously distributed in the spinal cord of vertebrate fish, and any fragments of the spinal cord are able to generate “fictive” swimming patterns under the condition of treatment with glutamatergic activators including NMDA, kainite, L-Glu and D-Glu (e.g., Poon, 1980; Dale and Roberts, 1984; Cangiano and Grillner, 2005; Wiggin et al., 2012). In order to assess the possible ubiquity of CPG circuits for swimming in *Ciona* larvae, we segmented larval bodies into two pieces at various position along the longitudinal axis and examined movement patterns of the anterior and posterior fragments (Fig. 1). When the cut sites were located within the trunk, the anterior fragments, naturally, showed no movement [Fig. 1A; MI = 0, 19 cases in 19 trials (19/19)]. In cases with the cut sites being located posterior to 5.7%, in which the tail length was regarded as 100%, the tail remaining in the anterior fragments exhibited evident alternating beats (Fig. 1A, Movie 1; MI = 4, 100/100). In cases with the remaining tail being shorter than 5.7%, the anterior fragments were silent (Fig. 1A; MI = 0, 6/6). These observations suggest that the proximal 5.7% portion of the tail in combination with the trunk is necessary to express locomotor output with an alternating pattern and that the distal 94.3% of the tail is not required for generating patterned locomotion.

On the other hand, posterior fragments did not show any evident movements when the fragments were 45% or shorter from the posterior end of the tail (Fig. 1B; MI = 0, 22/22). When posterior fragments were longer than 45%, if they were composed only of the tail (namely the cutting sites were in 0∼55%), they exhibited sporadic and irregular movements with higher frequency (≥ 3 Hz; Movie 1; MI = 2, 33/84) or lower frequency (< 3 Hz, MI = 1, 47/84). Almost all of these movements observed were not regularly patterned (only one exception in 84 trials) (Fig. 1B). However, posterior fragments containing a portion of the trunk, namely the cut sites being in the trunk, exhibited patterned locomotor output (Fig. 1B, Movie 1; MI = 4, 18/19). Considering that a posterior fragment cut at -6.7% represented the only case in which clear patterned tail beatings were not observed in our series of experiments, it can be concluded that a posterior fragment containing posterior -6.7∼0% or more of the trunk sufficiently generates patterned output as the longest estimation and therefore the anterior portion of the trunk to this level is not necessary. The results also revealed that a reciprocal tail beating pattern is not generated by body parts composed only of the tail region (105/106 cases), and thus a small portion of the posterior trunk (-6.7∼0% or less) is required for generating patterned movements.

It should be noted that all of the results shown above were obtained from body fragments just placed in ASW without any additional stimuli. Previously, Nishino et al. (2010) reported that swimming movements of *Ciona* larvae in which a large anterior portion of the trunk had been removed could be activated by L-Glu treatment, but our present results showed that locomotor output in decapitated *Ciona* larvae can be sufficiently driven without any activator agonists. We performed the experiment described above in ASW containing 100 μM L-Glu, but region-specificity of the relationships between cut sites and movement patterns did not considerably differ from those without L-Glu (Fig. S1).

### Mid-piece fragments sufficiently express patterned movements

The results presented above suggested that the middle part of the larval body (herein called “mid-piece”) spanning -6.7∼5.7%, as the longest estimation, represents a necessary and sufficient region to generate reciprocal tail beating pattern. In fact, mid-piece preparations that lacked most of the anterior part of the trunk and also most of the posterior tail clearly expressed alternating patterned tail beatings (Movie 2; MI = 4, - 6.5∼19.2% mid-piece in the indicated case). To assess the quantitative relationships between the remaining part/length of mid-piece fragments and frequency of alternating tail beats, we analyzed the movement patterns generated by the mid-piece fragments using a high-speed camera. This quantitative analysis revealed that the mid-piece fragments generate alternating tail beats of 10.4 ± 1.4 Hz (mean ± s.d.; n = 22), whose levels appeared irrespective of the remaining part/length of the fragment (Fig. 2). We examined again the effect of L-Glu (100 μM) on the frequency of tail beating expressed by mid-piece fragments, but the frequency did not significantly differ from that by untreated mid-piece fragments (10.4 ± 1.8 Hz in 100 μM L-Glu, n = 9; p > 0.1, Student’s *t*-test, two-tailed). We did not find any evident dose-dependent increase of swimming activities, average duration of tail beating bursts or cumulative duration of bursts per minute in ASW containing 0, 100 μM, or 1 mM L-Glu (Fig. 3; p > 0.05 in every combination, Student’s *t*-test, two-tailed).

**Fig. 2.**
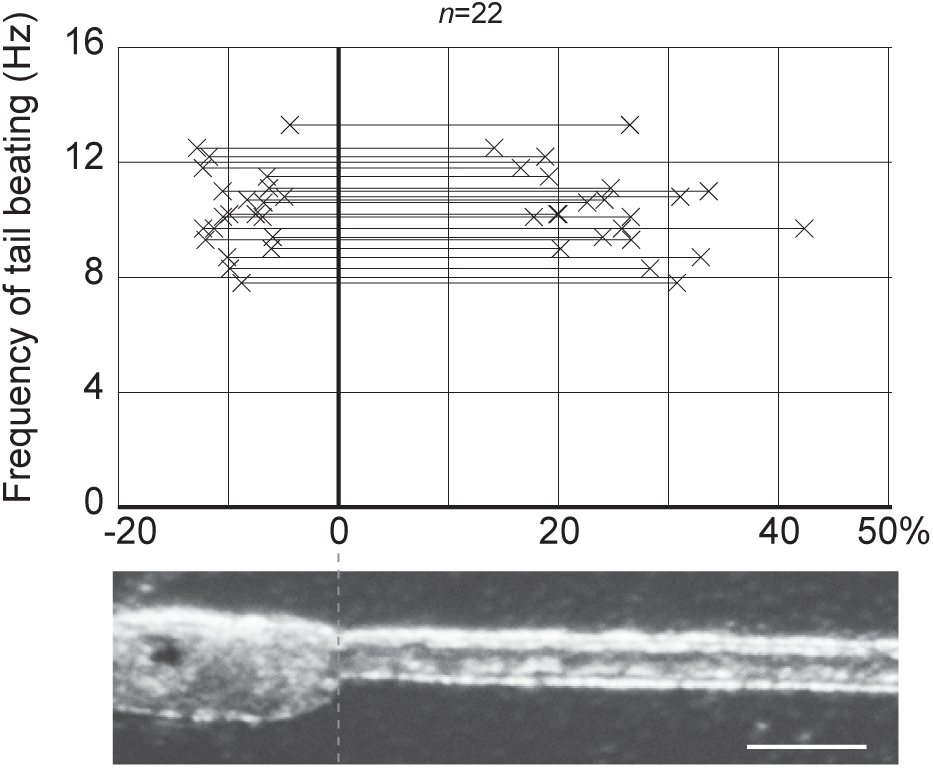
Frequency of tail beating in “mid-piece” fragments of *Ciona* larvae. Mid- piece fragments from which the anterior trunk and posterior tail have been excluded show 8-12 Hz tail beating. Cut sites (marked by ×) are indicated along the horizontal axis, which is defined to be 0% and 100% at the trunk-tail junction and at the tail tip, respectively. Horizontal bars between the pairs of ×’s indicate the remaining portion of the body in the mid-piece fragments. Picture of a *Ciona* larva is shown for reference. *n* denotes the number of fragments examined. Scale bar = 100 μm.

**Fig. 3.**
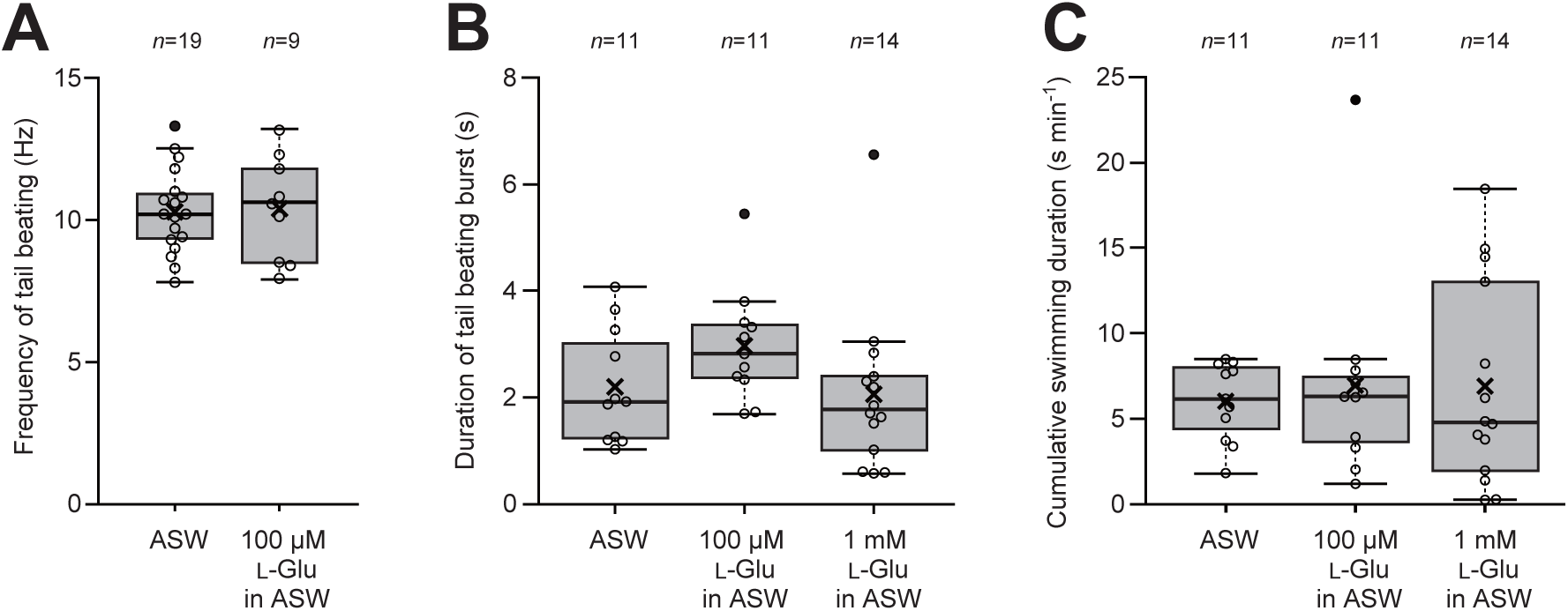
Tail beating activity of mid-piece fragments in the presence of L-Glu. (A) Frequencies of tail beating exhibited by mid-piece fragments in the absence of L-Glu (ASW) and in the presence of L-Glu (ASW with 100 μM with L-Glu). (B) Durations of tail beating burst in the absence of L-Glu (ASW) and presence of L-Glu (ASW with 100 μM or 1 mM L-Glu). (C) Cumulative durations of tail beating bursts in one minute (length of time in which mid-piece fragments express tail beatings during one minute), in the absence of L-Glu (ASW) and presence of L-Glu (ASW with 100 μM or 1 mM L-Glu). *n* denotes the number of fragments examined. Open circles indicate data utilized for boxplots, and filled circles indicate outliers. “×” indicates mean values.

### Mid-piece of *Ciona* larva exhibits periodicity of patterned tail beatings

In the course of examination of locomotion patterns of mid-piece fragments, we noticed that these fragments expressed swimming bursts with almost regular periodicity (Fig. 4, Movie 3). We dissected 22-24 hpf swimming larvae to isolate mid-piece fragments and measured the lengths of time from the beginning of a swimming event to that of the next event for a period of 20 min after amputation. Our observations clearly demonstrated intervals of approximately 20 s between the bursting events (Fig. 5; in 12/16 cases, 50% or more of the events were included in the neighboring top 3 ranks with the peak in 8-26 s). In a general meaning, random events that occur on a time series can be regarded as a Poisson arrival process (Fig. 5A). In such a case, the distribution of time intervals between events should be on a negative exponential curve (λe^-λt^), where the time constant (= λ^−1^ = τ) represents the mean value of intervals as well as the standard deviation (s.d.). If swimming burst events expressed by mid-piece fragments were in a random process, a histogram of lengths of time between the events would be close to the negative exponential distribution with a time constant ≈ 20 s, but the distribution of interval data collected in reality appeared much different from such a random process [Fig. 5A; cf. histograms of lengths of time between the tail beating bursts expressed by a mid-piece fragment, where τ = 17.5 s (dark gray), and of intervals of randomly occurring events (calculated, light gray)]. There was only one case (1/15) that showed a negative exponential pattern of distribution with the mean value close to the s.d. value (Fig. 5B, no. 8).

**Fig. 4.**
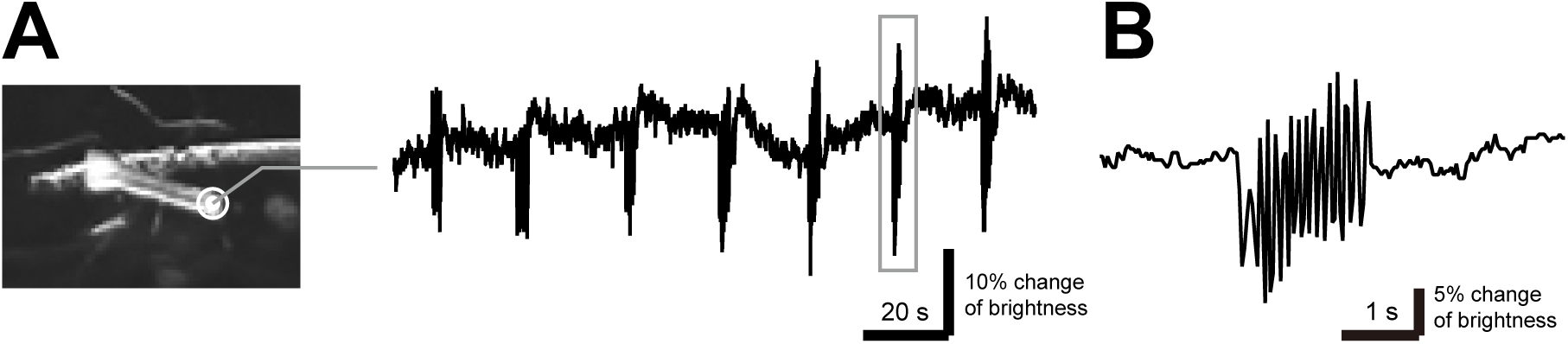
Periodicity of tail beating bursts expressed by mid-piece fragments. (A) Point of interest (double circle) was set on the posterior tip of a mid-piece fragment (left panel). Brightness changes at that point that reflect alternating tail beats (right panel). Fluctuation of baseline reflects subtle changes in the position of the fragment as well as noise in a series of images, not beating of the tail. (B) Magnified trace of a tail beating burst (indicated by the rectangle in A). Alternating tail beats of ∼10 Hz were detected on the trace.

**Fig. 5.**
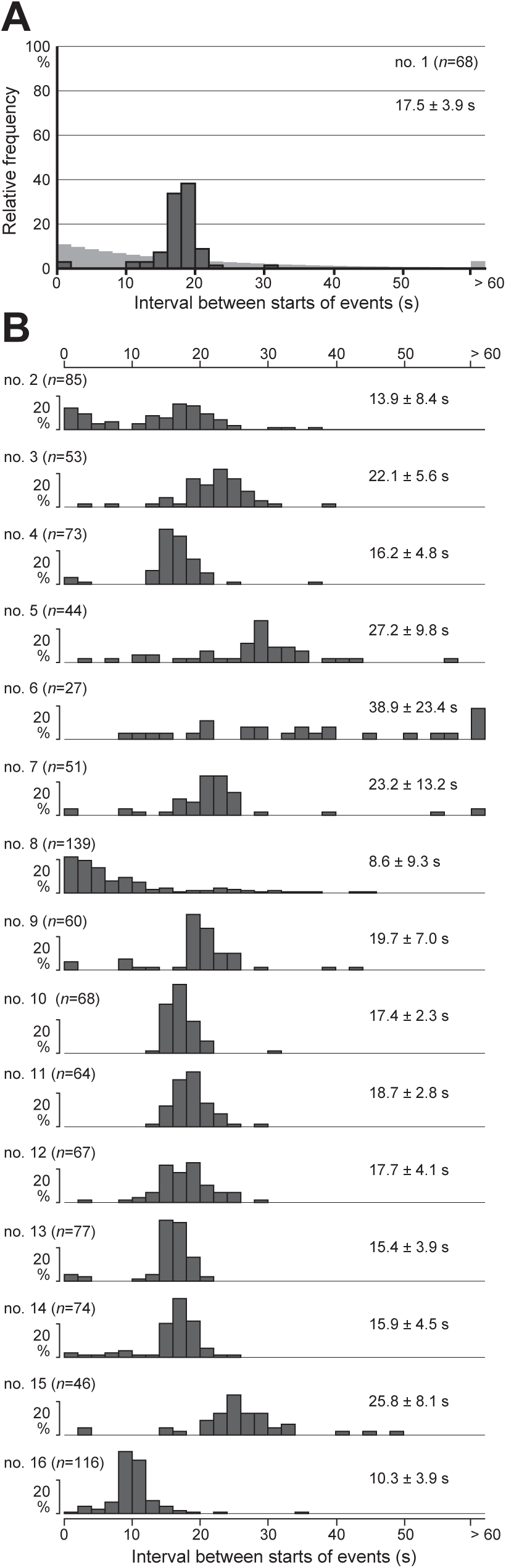
Cycle periods of tail beating bursts exhibited by mid-piece fragments. (A) Histogram showing cycle periods of tail beating bursts expressed by a mid-piece fragment (about 20 min after amputation) from the start time of one burst to the start time of the next burst (dark gray). Relative frequencies of lengths of cycle periods in 68 cycles (depicted by *n*) and their mean ± s.d. values are shown. Light gray bars are a reference to illustrate the distribution pattern of relative frequencies when bursts occurred randomly with the same average time between events (i.e., along “random arrival events”, see also text). (B) Histograms showing cycle periods of tail beating bursts expressed by 15 other mid-piece fragments (no. 2-16) (dark gray bars). *n* denotes the number of examined cycle periods of tail beating bursts. Mean ± s.d. values among lengths of cycle periods in each fragment are also shown. Vertical ticked axes represent 20% of relative frequency.

This periodic pattern of swimming events may be affected by amputations or other related damage. In order to estimate the degree to which these activities of mid- piece fragments were affected by our manipulations, the fragments were left for about 1.5 h to allow time for the fragments to recover from possible damage (or possibly to be deathly silent), and then it was examined whether they exhibit periodic bursts. We prepared mid-piece fragments derived from 22-24 hpf larvae, left them for 1.5 h or more, and then measured durations of patterned tail-beating events (ignoring sporadic flick-like twitches) (Fig. 6A, gray bars) as well as intervals of tail beating bursts (Fig. 6B, black bars). We measured durations (start-end) and intervals (end-start) of tail beating bursts in this analysis, not lengths of time between neighboring start-start timings as described above, since we consider that it would not be realistic for a PG to encode periodic “start time” but to keep the “interval” from the end of a burst to the start of the next burst to be even. Our observations revealed that periodic expressions of locomotor patterns are retained even at 1.5 h after amputations (Fig. 6A,B; in 13/17 cases, 50% or more of the intervals were included in the neighboring top 3 ranks with the peak in 10-30 s), and the duration and interval of tail beating bursts were estimated to be 2.4 ± 1.9 s and 19.8 ± 8.4 s, respectively [mean ± s.d.; N (individual no.) = 17, n (event no.) = 866]. This estimation showed that the cycle period became slightly longer than that immediately after the amputation (cf. Figs. 5B and 6A,B). This periodic pattern of locomotion was also observed for mid-piece fragments derived from earlier larvae (19-20 hpf) (Fig. S2; in 13/17 cases, 50% or more of the intervals were included in the neighboring top 3 ranks with the peak in 10-30 s), in which the duration and interval of tail beating bursts were 2.8 ± 3.0 s and 21.0 ± 8.1 s, respectively (N = 17, n = 781).

**Fig. 6.**
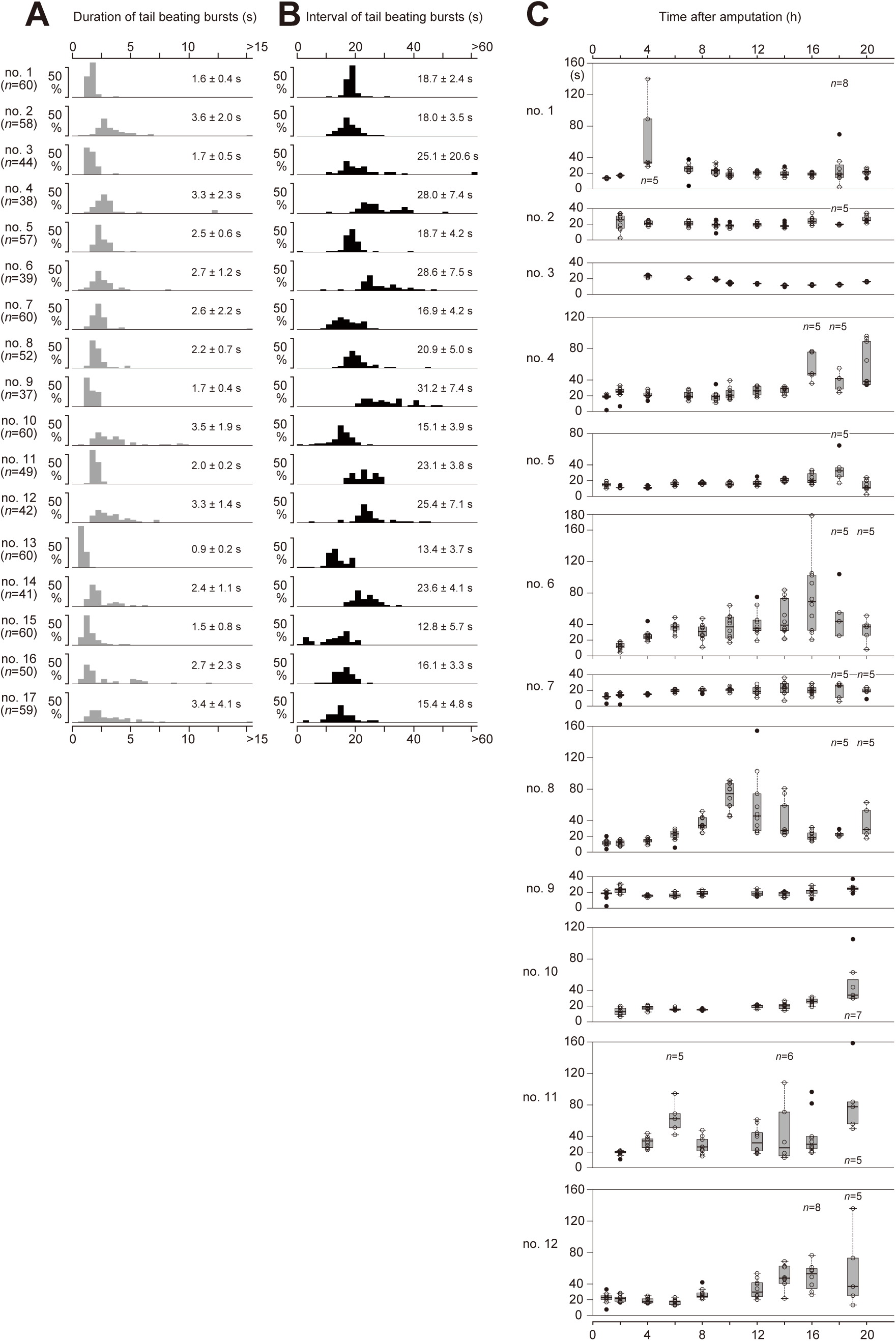
Stability of cycle periods of tail beating bursts expressed by mid-piece fragments. (A,B) Histograms showing durations (A) or intervals (B) of tail beating bursts expressed by mid-piece fragments (1.5 h after amputation at 22-24 hpf). *n* denotes the number of examined cycle periods of tail beating bursts in each specimen. Mean ± s.d. values of durations and intervals in each fragment are also shown. Vertical ticked axes represent 50% of relative frequency. (C) ∼20-s interval of tail beating bursts is maintained for a half day or longer. Intervals between 11 (or less) consecutive bursts expressed by mid-piece fragments (no. 1-12) were measured every 0.5-2.0 h up to 20 h after amputation at 22-24 hpf. Open circles indicate data utilized for boxplots, and filled circles are outliers. *n* represents the number of measured intervals; only in cases in which they are less than 10, *n*’s are indicated above or below the boxplots.

We further measured intervals of tail beating bursts up to 20 h after the amputations at 22-24 hpf, and we found that the majority of the prepared mid-piece fragments continued to show periodicity for 12 h or longer (Fig. 6C). It is of note that all the mid-piece fragments continued tail beatings without stopping the cycle or exhibiting metamorphosis for such a long time. These observations demonstrated that the mid-piece of *Ciona* larvae that possesses a property to perform alternating tail beats expresses an intrinsic, autonomous, and robust rhythm to generate ∼2.5-s tail beating bursts with ∼20-s intervals.

### Periodicity of swimming events in unfragmented larvae

Next, we examined how the periodicity observed for mid-piece fragments is represented in swimming patterns expressed by whole bodies of *Ciona* larvae. We considered that it would be difficult to keep tracing temporal patterns of a single larva that freely swims in seawater and also that it would be difficult to exclude accidental sensory stimulations on the larva such as uncontrolled physical contact with the wall and bottom of the container or the the water surface. Thus, we placed a larva in seawater containing methylcellulose (MC), which provided higher viscosity to the medium, in order to limit their translocation and keep their sensory world constant as much as possible.

We confirmed by using a high-speed camera that frequencies of tail beats were not so different among larvae (∼24 hpf) in ASW with 0%, 0.5%, 1%, and 2% MC (Fig. S3), although the addition of 2% MC slightly decreased the tail beat frequency. This analysis revealed that addition of 1% or less of MC and accompanying increase of viscosity did not greatly affect their swimming performance, while it was shown that the whole body of larvae exhibits a higher frequency of tail beats (15-20 Hz) than do mid- piece fragments (8-12 Hz) (cf. Figs. 2, 3A and Fig. S3).

In the presence of 1% MC, we observed unfragmented larvae (19-20 hpf) to measure durations and intervals of tail beating bursts. Durations of patterned tail beatings were greatly variable, but their intervals were not along a negative exponential curve, namely not random (Fig. 7A,B; in 9/17 cases, 50% or more of the intervals were included in the neighboring top 3 ranks with the peak in 8-26 s). Some periodicity in intervals was found to be shorter than 20 sec (Fig. 7B,E; 15.0 ± 9.5 s, N =17, n = 567). When we further decreased the physical stimulation on to larvae by using an optical filter (high-pass 590 nm) to eliminate the light visible to the *Ciona* larvae (Nakagawa et al., 1999; Nishino et al., 2011), the interval time between swimming bursts became even shorter (Fig. 7D,E; 10.4 ± 7.4 s, N = 10, n = 810) with higher periodicity [variance in the dark = 54.2 (n = 810) vs. variance in light = 90.3 (n= 567)]. On the other hand, variance in durations of the bursts by whole larvae placed in light was substantially higher than that by mid-piece fragments [Fig. 7E; variance in whole larvae in light = 168.9 (n = 567) vs. variance in mid-piece fragments = 9.2 (n = 781)], suggesting that efferent and afferent inputs, which are derived through the BV and other sensory systems into the mid-piece, have effects to shorten or lengthen swimming durations (Fig. 7A,E). In the dark, durations of tail beating bursts appeared almost randomized (Fig. 7C,E). These results suggest that *Ciona* larvae have a non-random temporal pattern in intervals of swimming bursts and that the larvae are able to modify, by means of parts other than the mid-piece, periodicity for swimming intervals and durations accommodated to sensory inputs.

**Fig. 7.**
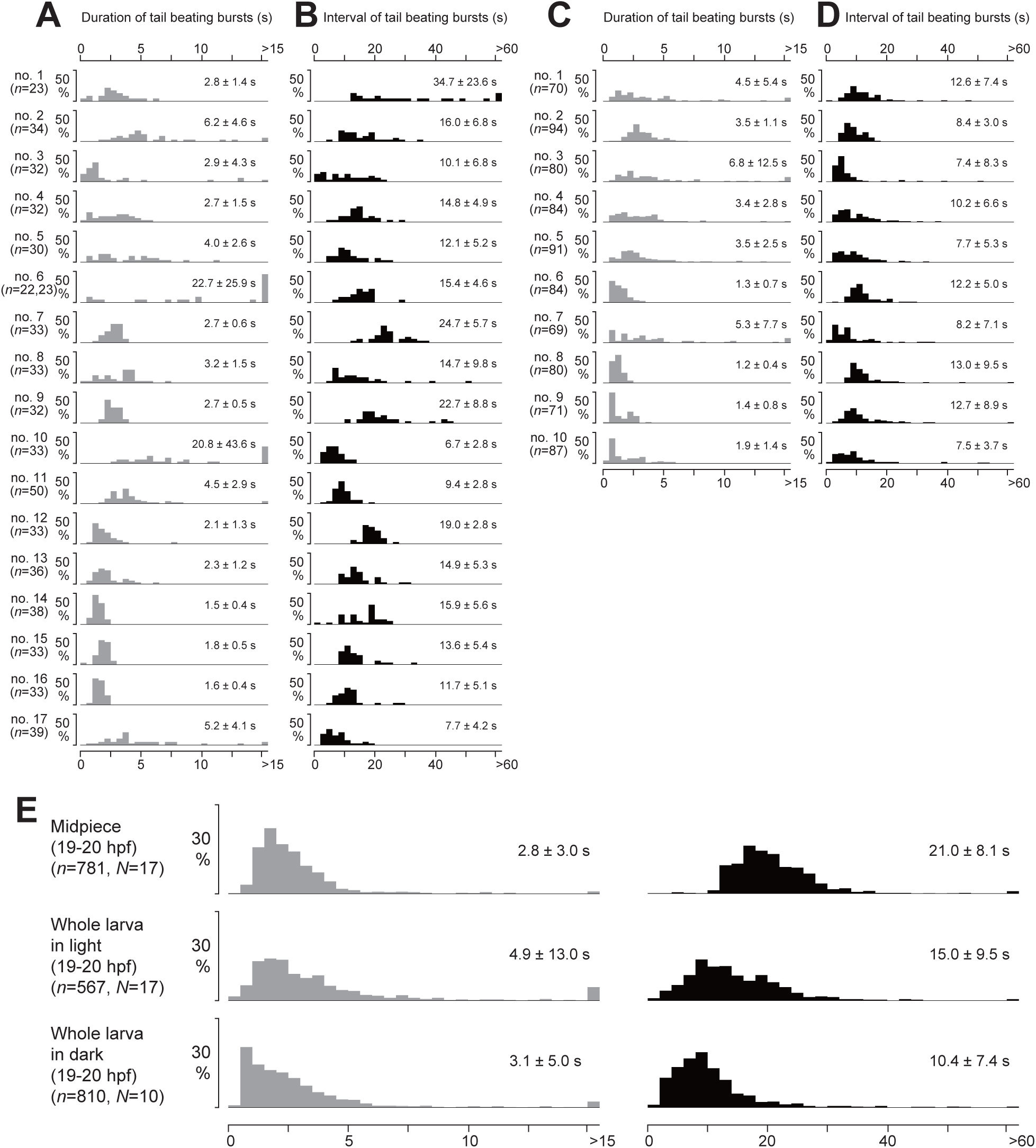
Temporal patterns of tail beating bursts expressed by unsegmented whole larvae in light and dark conditions. (A-D) Histograms showing durations (A,C) or intervals (B,D) of tail beating bursts expressed by unsegmented whole larvae (19-20 hpf) placed in 1% MC under a light (A,B) or dark (C,D) condition. *n* denotes the number of examined cycle periods of tail beating bursts in each specimen. Mean ± s.d. values of durations and intervals in each fragment are also shown. Vertical ticked axes represent 50% of relative frequency. (E) Histograms of aggregated data from different conditions. Total incidences of durations (left, gray bars) and intervals (right, black bars) of tail beating bursts expressed by mid-piece fragments amputated at 19-20 hpf (top, N = 17; corresponding to the data shown in Fig. S2A,B), whole larvae (19-20 hpf) placed in a light condition (middle, N = 17; corresponding to those shown in A,B), and whole larvae (19-20 hpf) placed in a dark condition (bottom; corresponding to those shown in C,D) are shown. *n* denotes the summed number of cycle periods. Mean ± s.d. values of durations and intervals are also shown. Vertical ticked axes represent 30% of relative frequency.

## DISCUSSION

### A swimming PG for *Ciona* larvae is localized to the mid-piece region

In this study, we identified a body region that is required for and is sufficient for expressing alternating tail beating bursts. The body region includes a least part of the posterior trunk and approximately 6% of the anterior portion of the tail, representing one-tenth or less of the length of the ∼1 mm whole body. Other regions did not show sufficient ability to express patterned movements in the absence or presence of an activation amino acid, L-Glu, in the present study. We can conclude that the posterior region of the tail is never sufficient for patterned swimming. On the other hand, we cannot determine if the anterior region of the trunk is insufficient since it essentially lacks an effector to express locomotor output. Therefore, we conclude here that most of the trunk excluding the posterior least portion is not required for generating patterned movements. Our results show the distinction of this system found in the *Ciona* larva from what has been uncovered in the spinal cord of vertebrate fish or amphibian tadpoles, in which CPG neuronal circuits are not localized but are ubiquitously distributed along the spinal cord (e.g., Roberts, 2000; Wiggin et al., 2012; Grillner and El Manira, 2020).

We found that posterior fragments that do not include the posterior trunk could indeed move frequently but did not show patterned tail beats with rhythmic alternations (only one exception) (Fig. 1B). This clearly indicates that the posterior portion of the trunk is required to express patterned movements. In contrast, short posterior fragments (segmented at 55-100%) did not show any movements. Considering that muscle cells occupy up to ∼85% length of a *Ciona* larval tail (see Nishino et al., 2011), it would be natural that posterior fragments of ∼15% (85∼100%) or shorter from the tail tip do not move. However, it is intriguing why posterior fragments segmented between 55% and 85% did not move, whereas most of longer posterior fragments cut at 55% or less moved. Irreversible damage on to muscle cells near the cut site seems not feasible; otherwise anterior fragments with a short remaining tail would not move either. One possibility is that the midtail neurons (MTNs) that are sparsely distributed along the CNC and express the vesicular ACh transporter, a representative marker for cholinergic neurons, are present in the longer posterior fragments but not in the shorter ones (referred to as planate cells in Imai and Meinertzhagen, 2007; see also Horie et al., 2010; Ryan et al., 2016). Since larval muscle cells are activated by cholinergic input (Nishino et al., 2011), MTNs may constitute a group of motor neurons, and their spontaneous excitations may underlie the sporadic movements of longer posterior tail fragments. Larger number of MTNs would be included in longer posterior fragments of the tail; longer tail fragments actually tended to move more frequently. Two pairs of bipolar tail neurons (BTNs) are also known to reside in the middle of the tail (Coric et al., 2008; Stolfi et al., 2015), but they express the enzyme for synthesizing an inhibitory neurotransmitter GABA (Zega et al., 2008; Stolfi et al., 2015). If BTNs are GABAergic in reality, it would not be easy to attribute their movements to spontaneous firing of BTNs.

Anterior fragments expressed patterned tail beats when the fragments contained at least 5.7% of the proximal portion of the tail. In this proximal tail portion, there are two pairs of contralateral inhibitory neurons called ACINs (Horie et al., 2010; see also Nishino et al., 2010). The results of the present study provide direct evidence for the requirement of ACINs to generate alternating tail beats, which has long been proposed (Horie et al., 2010; Nishino et al., 2010). In this study, we originally assumed the possible occurrence of unilateral rhythmic movements in some fragments and made “rank 3” in our motion index as to represent a type of patterned movements (Table 1). All of the patterned movements observed in larval fragments, however, appeared alternating, and no examples with unilateral rhythmic contractions of the tail (MI = 3) were found in our trials (> 200). Our observations support the idea that interaction between the left and right halves enables reciprocal movements. Indeed, the anterior pair of ACINs seems to be included in the proximal 5% of a *Ciona* larval tail (see Horie et al., 2010), suggesting that at least one pair of ACINs is required. However, considering slight shrinkage of our bisected fragments as mentioned above (see Methods) and possible variation of antero-posterior locations of the neuron types among individuals, it is not appropriate for us to conclude functional difference between the anterior and posterior pairs of ACINs. On the other hand, the reason why longer amputation of the posterior tail (cutting at 0∼5.7%), namely total elimination of ACINs, led to complete loss of movements, not to irregularity of movements, is unknown. One simple explanation is a physical limitation that movements of fragments with very short portions of the tail could not be detected by our eyes. Despite these ambiguous issues remaining, it is valuable to indicate that the data obtained here from a series of simple amputations of larvae clearly uncovered functional regionalization to express locomotor outputs. This in reverse implies that the distribution pattern of functional units, especially of endogenous neuron types, along the anterior-posterior axis is not so much varied among *Ciona* larval individuals (see Ryan et al., 2016; Nishino, 2018).

The CPG is, by definition, neural circuits in the CNS to express rhythmic motor outputs without any sensory inputs or feedbacks (Bässler, 1986; Delcomyn, 1998). In this meaning, it may still be precocious to claim that our “mid-piece” preparation represents a CPG that is free of sensory inputs. By amputating large portions of the anterior trunk as well as the posterior tail, most of the sensory neurons that have been identified so far, including papillar neurons, most of the ENs, and BTNs, would be excluded, but some ENs, such as some of the pACENs, DCENs and VCENs, might remain (Yokoyama et al., 2014; Ryan et al., 2018). Afferent inputs from ENs into the CNS tend to concentrate in the posterior part of the BV, e.g., Eminens cells (Horie et al., 2008a; Ryan et al., 2018), and other dense inputs are also found in the dorsal area of the MG, where AMG neurons reside (Ryan et al., 2018). Although ENs in general express vesicular glutamate transporter, a maker for glutamatergic neurons (Horie et al., 2008a), our results showing that L-Glu treatment did not change movement patterns of the mid- piece fragments suggest that inputs from ENs possibly remaining therein did not have much influence on the generation of tail beating patterns. We therefore propose that there is a neural circuitry that sufficiently fulfils the criteria for defining a CPG restrictedly and sufficiently present in the region of the trunk-tail junction but absent in the posterior ∼94% region of the tail (Fig. 8).

**Fig. 8.**
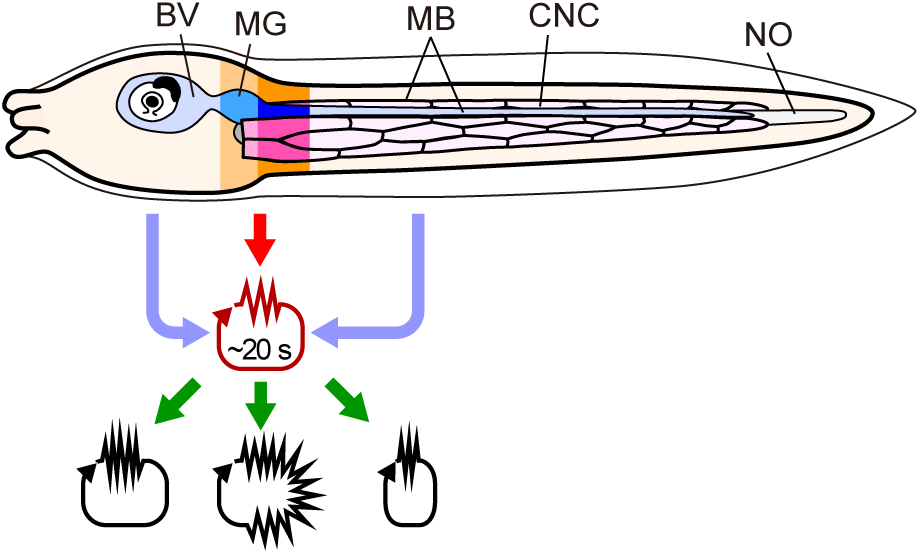
A proposed model for expression of swimming performance in the *Ciona* larva. The region highlighted with darker colors indicates the required “mid-piece” part for expressing autonomous reciprocal tail beating. The slightly darker part indicates a largest estimation, and the much darker part highlights the predicted region containing the sufficient PG. This mid-piece part harbors potential to autonomously output tail beating bursts with a ∼20-s cycle period. Other regions possessing various sensory apparatuses, such as photoreceptors and a gravity sensor (represented by pigments in the BV), modify this endogenous and autonomous drive to form swimming performance, which is accommodated to the sensory inputs. BV, brain vesicle; CNC, caudal nerve cord; MB, muscle band; MG, motor ganglion; NO, notochord.

### The mid-piece of the *Ciona* larva expresses an autonomous cycle period of tail beating bursts

The mid-piece fragment has the ability to express autonomously and periodically a locomotor pattern. By referring to the results of connectome analyses reported by Ryan et al. (2016, 2017, 2018), we could estimate that this restricted region in the mid-piece contains 25 neurons, at most, in the MG and 1 or 2 pairs of ACINs posteriorly associated to the MG (27-29 neurons in total) in the CNS. Five pairs of motor neurons that extend axons posteriorly along the CNC have been identified in the MG, which is located at the posterior dorsal edge of the trunk (Katz, 1983; Ryan et al., 2016, 2017). Considering our results showing that a small portion of the posterior trunk sufficiently confers ability to generate patterned movements, a posterior portion of the MG containing posterior pairs of motor neurons, not all of them, would constitute a sufficient circuit (Fig. 8, darker area). We observed only one case in which a posterior fragment cut within the tail (at 2.5%) exhibited patterned movements, which we consider is probably because of a slight extension of the posterior part of the MG into the tail in this larva and/or of excessive deformation of fragments leading to some error in determining the cut site.

How the neuronal network in the posterior MG and ACINs autonomously expresses alternating tail beats with a ∼20-s cycle period is an interesting question. In mammalian brain-slice preparations, “autonomous” rhythm-generating neural circuits have been found, including motor pattern/rhythm centers for breathing such as medullary preBötzinger and Bötzinger complexes (Smith et al., 1991; Koshiya and Smith, 1999; Ghali, 2019). Anesthetization makes respiration rhythms slower, but rhythmic respiration is kept at a “basal” level, indicating that the basal rhythm for breathing is autonomous in these complexes and also that it can be upregulated by various modifiers such as unvoluntary homeostatic feedbacks (O2 concentration, body temperature, etc.) and intentional controls (sighing, etc.). Paralleling these, the mid-piece fragment of *Ciona* larvae expressed slower tail beatings and longer (but more periodic) intervals between their bursts, representing a basal status, and this basal rhythm was upregulated (i.e., tail-beat frequency elevated and burst intervals shortened) by inputs from other parts (see Fig. 3A, 7E, and S3). The rhythm generator found in the *Ciona* mid-piece and the mammalian preBötzinger complex share intrinsic properties to autonomously express rhythmic bursts and to be upregulated by signals from other parts, although the mid-piece of the *Ciona* larva outputs a left-right reciprocal pattern. We did not obtain any idea from our results on what kinds of neurons in the mid-piece fragment generate autonomous rhythm and how they do that. Recently, Akahoshi et al. examined firing patterns of embryonic cells including differentiating neurons using a fluorescent Ca^2+^ indicator (Akahoshi et al., 2017), and they very recently reported that a pair of neurons positioned in the posterior MG autonomously elevate intracellular Ca^2+^ concentration that corresponds to tail beatings with ∼20-s intervals in the late tailbud stage (Akahoshi et al., 2021 preprint). The oscillatory rhythm they found in the posterior MG corresponds well with the rhythm of swimming bursts found in our mid-piece preparations; that pair of neurons may be an origin of the swimming rhythm generation.

### Body regions outside the mid-piece modify its autonomous activity of the mid-piece

Ascidian larvae intermittently execute tail beating to swim, and the larvae gravitate downward during intervals. Considering that the larvae have a tendency to exhibit negative gravitaxis (Svane and Young, 1989; Tsuda et al., 2003a; Bostwick et al., 2020), temporal control of durations and intervals of larval tail beating would be a crucial strategy for sessile ascidians to disperse progeny. Indeed, duration time of tail beating bursts was reported to be under the control of neurotransmission (Brown et al., 2005). Based on the results of our study, we propose that swimming performance expressed by the whole *Ciona* larva is represented through modifications to a slower basal rhythm intrinsic to the mid-piece circuitry by multiple input signals from other parts of the body (Fig. 8). Sporadic movements, for example flicks, of the tail can autonomically, but irregularly, be generated by machineries residing in the posterior 6∼100% region of the tail, which may also stimulate the mid-piece circuitry in turn. These provide a mechanistic basis for temporal control of swimming pattern generation in *Ciona* larvae (Fig. 8).

Body regions outside the mid-piece had effects to increase variation in the duration of swimming bursts and to shorten intervals of the bursts. Moreover, in larvae with the BV placed in a dark condition, swimming intervals became further shortened. Regarding efferent neural signals from the photoreceptors in the BV to MG neurons, both excitatory and inhibitory visuomotor pathways have been indicated so far (Kourakis et al., 2019; Bostwick et al., 2020). Therefore, our results shown in Fig. 7 would reflect elevated excitatory and/or suppressed inhibitory visuomotor signals in the constant dark condition (Kourakis et al., 2019). The unsegmented larvae used in this study were young larvae not long after hatching (19-20 hpf). *Ciona* larvae are known to gradually develop and mature in various aspects such as the ability to respond to light stimuli (Kajiwara and Yoshida, 1985; Tsuda et al., 2003b; Zega et al., 2006; Horie et al., 2008b; Salas et al., 2018). If larvae at different developmental times had been used, different patterns of modifications on to the mid-piece, the properties of which are relatively constant, might have been observed. This point remains to be clarified.

Swimming motor patterns of vertebrates are expressed by ubiquitously distributed CPGs along the spinal cord, and CPGs are reciprocally inhibited between the left and right sides (e.g., Roberts et al., 2010; Grillner and El Manira, 2020). These neural circuits in the spinal cord are thought to be basically silent without an activating signal and are driven in vivo by descending projections from locomotor regions in the diencephalon and mesencephalon (for review, Grillner and El Manira, 2020). Namely, locomotion driven by the spinal cord of vertebrate swimmers is basically under the control of voluntary inputs from superior control centers in the brain. In tonically activated states of the spinal cord (e.g., by bath application of N-methyl-D-aspartate), durations of swimming bursts as well as intervals between the bursts become shorter in a dose-dependent manner, suggesting that the amount of stimulus not only affects burst frequency but also changes the timing of ON and OFF of the bursts (Cangiano and Grillner, 2003; Wiggin et al., 2012). The motor circuit in the mid-piece of *Ciona* larvae has a property to express alternating and periodic bursts, which is reminiscent of that of CPGs in the spinal cord of swimming vertebrates (Grillner, 1975; Wiggin et al., 2012; Grillner and El Manira, 2020). On the other hand, the *Ciona* larval motor pattern is also reminiscent of breathing motor patterns, in that it is based on an autonomous, involuntary, and slower rhythm of bursts that secures the basal level of locomotion. This basic motor pattern is modified by several lines of inputs, which allows the larvae to swim in ad hoc ways in response to various sensory stimuli (even though we would not regard their swimming simply as “voluntary” or “intentional”). Ascidian larvae share tadpole-shaped bodyplans with vertebrates, including the axial notochord, dorsal nerve tube, and bilateral muscle bands, but the results presented here showed that there are several significant differences between the ascidian larva and vertebrate swimmers in the localized/ubiquitous presence of the CPG, minimal/vast redundancy of CPG circuits and, importantly, autonomy/non-autonomy of their activity. This central pattern/rhythm generator that facilitates *Ciona* larval locomotion represents the simplest one for chordate swimmers, one which is moreover to be resolved in identifiable cells (Ryan et al., 2016; Nishino, 2018; Gibboney et al., 2020; Akahoshi et al., 2021 preprint).

## Acknowledgments

We thank Drs. Satoe Aratake, Megumi Kotsuka, and Manabu Yoshida (Misaki Marine Biological Station, The University of Tokyo), Drs. Reiko Yoshida, Chikako Imaizumi, and Yutaka Satou (Laboratory of Developmental Genomics, Department of Zoology, Graduate School of Science, Kyoto University), and other staff who distribute *Ciona* under the National BioResource Project, AMED, Japan. We are grateful to Dr. Kohji Hotta (Keio University) for his discussions and for kindly sharing unpublished data. We deeply thank Dr. Yasushi Okamura and Dr. Ian A. Meinertzhagen for encouragement and critical reading of the manuscript.

## Competing interests

The authors declare no competing or financial interests.

## Author contributions

ASN conceived the research concept. ASN, TH and SH designed experiments, and TH and SH performed the experiments and data acquisition. YI substantially contributed to the establishment of a technical pipeline for analyzing the movements of larvae (or fragments) that had been recorded on movie files. ASN wrote the draft, and TH, SH, YI and ASN edited and approved the manuscript.

## Funding

This research was mainly supported by JSPS KAKENHI to ASN (No. 25440150, 17K19369, and 20K06713) and in part by Grants-in-Aid from Hirosaki University to ASN (Hirosaki University Grant for Exploratory Research by Young Scientists during 2012-2015 and 2017, Hirosaki University Institutional Research Grant for Young Investigators 2017-2019, and Interdisciplinary Collaborative Research Grant for Young Scientists, Hirosaki University in 2018 and 2019) and also in part by The Yamada Science Foundation, The Sumitomo Foundation, and The Sekisui Integrated Research Foundation to ASN.

## Notes

### Competing Interest Statement

The authors have declared no competing interest.

